# Interfering with MIF-CD74 signalling on macrophages and dendritic cells with a peptide-based approach restores the immune response against metastatic melanoma

**DOI:** 10.1101/248807

**Authors:** Carlos R. Figueiredo, Ricardo A. Azevedo, Sasha Mousdell, Pedro T. Resende-Lara, Lucy Ireland, Almudena Santos, Natalia Girola, Rodrigo L.O.R. Cunha, Michael C. Schmid, Luciano Polonelli, Luiz R. Travassos, Ainhoa Mielgo

**Affiliations:** Department of Molecular and Clinical Cancer Medicine, University of Liverpool, Liverpool, United Kingdom.; Experimental Oncology Unit (UNONEX), Department of Microbiology, Immunology and Parasitology, Federal University of São Paulo (UNIFESP), São Paulo, SP - Brazil.; Laboratory of Computational Biology and Bioinformatics, Federal University of ABC, Santo André, São Paulo, SP, Brazil.; Laboratoire de Biologie et Pharmacologie Appliquées (LBPA), UMR 8113, Ecole Normale Supérieure, Cachan, France.; Chemical Biology Laboratory, Natural and Human Sciences Center, Federal University of ABC, Santo André, SP, Brazil.; Unit of Biomedical, Biotechnological and Translational Sciences, Department of Medicine and Surgery, Universitá degli Studi di Parma, Parma, Italy.

**Author notes:** Senior corresponding author: Dr Ainhoa Mielgo, Department of Molecular & Clinical Cancer Medicine, Institute of Translational Medicine, First Floor Sherrington Building, Ashton street, Liverpool L69 3GE, United Kingdom. Co-corresponding author: Dr Luiz R Travassos Department of Microbiology, Immunology and Parasitology, Federal University of São Paulo (UNIFESP), Vila Clementino – CEP 04023-062, 8th floor. São Paulo, SP - Brazil (ext: 1516 and 1517).

**Keywords:** metastatic melanoma, macrophages, dendritic cells, immune response, peptide-based immunotherapy, MIF, CD74.

## Abstract

Mounting an effective immune response against cancer requires the activation of innate and adaptive immune cells. Metastatic melanoma is the most aggressive form of skin cancer. Immunotherapies that boost the activity of effector T cells have shown a remarkable success in melanoma treatment. Patients, however, can develop resistance to such therapies by mechanisms that include the establishment of an immune suppressive tumour microenvironment. Understanding how metastatic melanoma cells suppress the immune system is vital to develop effective immunotherapies against this disease. In this study, we find that the innate immune cells, macrophages and dendritic cells are suppressed in metastatic melanoma. The Ig-CDR-based peptide C36L1 is able to restore macrophages and dendritic cells’ immunogenic functions and to inhibit metastatic growth *in vivo*. Mechanistically, we found that C36L1 interferes with the MIF-CD74 tumour-innate immune cells immunosuppressive signalling pathway and thereby restores an effective anti-tumour immune response. C36L1 directly binds to CD74 on macrophages and dendritic cells, disturbing CD74 structural dynamics and inhibiting MIF signalling through CD74. Our findings suggest that interfering with MIF-CD74 immunosuppressive signalling in macrophages and dendritic cells using peptide-based immunotherapy can restore the anti-tumour immune response in metastatic melanoma. Our study provides the rationale for further development of peptide-based therapies to restore the anti-tumour immune response.

## INTRODUCTION

Cutaneous melanoma is a cancer that develops from melanocytes generally located in the epidermal basal cell layer of the skin. At very-early stages, single skin lesions can be promptly excised and the 5-year survival rate of melanoma is 98%. Beyond these stages, however, melanoma can metastasize to distant organs including lungs, liver, bones and brain, and the 5-year survival rate in stage IV drastically decreases to 15-20% (1, 2). The aggressiveness of melanoma is associated with a strong burden of somatic mutations (3), with different neoepitopes making melanoma cells immunogenic and boosting the immune response (4, 5). In order to evade the immune response, melanomas often activate negative immune checkpoint regulators (ICRs) such as PD-1 and PD-L1 or CTLA-4 that inhibit effector T cell and function in peripheral tissues or lymph nodes, respectively (6, 7). Inhibition of the immune checkpoint regulators with anti-PD-1 and anti-CTLA-4 antibodies enables T-cell-mediated killing of melanoma cells and significantly improved patient outcomes in recent years (5). However, immune checkpoint inhibitors (ICI) are only effective if effector T cells infiltrate the tumour. The generation of effector T cells requires the activation and function of antigen presenting cells (APCs), such as dendritic cells (DCs) and macrophages (8, 9). DCs and macrophages are cells from the innate immune system that are essential for starting and shaping the immune response against any damaged tissue, including cancer (7,10).

Tumour associated macrophages (TAMs) are one of the most predominant immune cells in melanomas, and the number of TAMs inversely correlates with patients’ outcome, in both early and late stages of melanoma (11). Macrophages can be polarised into M1-like anti-tumorigenic and M2-like immunosuppressive macrophages (12). We, and others, have shown that, in tumours, macrophages are often polarised into M2-like macrophages that support tumour cell proliferation, survival, metastasis, resistance to therapy, and suppress the antitumour immune response (12-16). Similarly, DCs can also acquire immunogenic or tolerogenic behaviours depending on their maturation status (17). Immunogenic DCs are usually fully mature and express costimulatory and antigen presenting molecules such as CD40, CD80, CD86 and MHC-II to support T cell activation and function (17, 18). Immunogenic DCs, however, may switch to a tolerogenic phenotype during cancer progression, which inhibits the activation and function of effector T cells (19, 20). Tumour cells often secrete factors that polarise macrophages into M2-like immunosuppressive macrophages and suppress DCs immunogenic functions leading to an immuno-suppressed tumour microenvironment (7, 16, 21). Thus, the understanding of how metastatic melanoma suppresses the immune system and developing therapies that restore a pro-inflammatory and immunogenic environment are important areas of research.

Bioactive peptides based on immunoglobulin complementary determining regions (CDRs) are promising candidates for adjuvant cancer therapy and can stimulate the innate immune system (22-24). We have previously shown that different CDR peptides display anti-tumour activities against melanoma, and are able to regulate receptors and transcription factors on both tumour cells and immune cells (24-28). Recently, we identified the C36 V_L_ CDR**-1** peptide (C36L1) as an anti-tumour CDR-based peptide that inhibits metastatic melanoma cells proliferation and growth *in vitro* and *in vivo* (24, 25). However, the mechanism by which C36L1 inhibits metastatic melanoma progression in a syngeneic model remains unknown.

In this study, we found that C36L1 inhibits metastatic melanoma only in mice that have a competent immune system. C36L1 supports M1-like anti-tumorigenic macrophages and restores DCs pro-inflammatory phenotype and immunogenic function. C36L1 activation of macrophages and DCs results in a significant increase in the infiltration of effector T cells in the metastatic lungs, leading to a marked decrease in the tumour burden.

Macrophage migration inhibitory factor (MIF) is highly expressed in metastatic melanoma lesions and is secreted by B16F10 metastatic melanoma cells. Previous studies have shown that MIF can induce an immunosuppressive environment that supports melanoma progression (29, 30). The pharmacological effects of blocking MIF in the context of metastatic melanoma are beginning to be investigated, and the mechanisms by which MIF suppresses the immune cells remain poorly understood. CD74 is MIF’s main receptor, and is highly expressed in macrophages and DCs (31, 32).

In the present study we show that the C36L1 peptide binds to CD74 in both macrophages and DCs, disturbing its structural dynamics and inhibiting the MIF-CD74 signalling and the immunosuppressive effect on macrophages and DCs. These findings highlight the MIF-CD74 axis as an important mechanism of macrophage and DC immunosuppression in metastatic melanoma, and provide a rationale for further evaluation of CDR-based peptides as therapeutic agents able to restore macrophages and DCs’ anti-tumour functions in metastatic melanoma.

## MATERIALS AND METHODS

### Peptides

Peptides were purchased from Peptide 2.0 (Chantilly, VA, USA). C36L1 peptide (KSSQSVFYSSNNKNYLA-NH2) and the irrelevant iCDR control peptide (CE48-H2, INSGGGGTYYADSVKG-NH2) were synthesized with an amide group in the C-terminus, at 95–98% purity, determined by High-performance liquid chromatography (HPLC) using a C18 column and subsequently analysed by mass spectrometry.

### Cell culture

Murine melanoma B16F10 cells were cultured in complete RPMI-1640 medium (Thermo Fisher, Waltham, MA, USA) supplemented with 10 mM N-2-hydroxyethylpiperazine-N2 ethane sulfonic acid (HEPES), 24 mM sodium bicarbonate, 40 mg/L gentamicin, pH 7.2 and 10% fetal bovine serum (FBS), at 37°C, under humid atmosphere and 5% CO_2_. Primary macrophages and primary myeloid DCs were generated from C57BL/6-mice bone-marrows and cultured in DMEM-Dulbecco's Modified Eagle Medium (Thermo Fisher) and RPMI-1640 medium, supplemented with 10% (v/v) FBS, 2 mM L-glutamine, and 1X penicillin/streptomycin at 37°C, respectively, under humid atmosphere and 5% CO2. Cells were regularly checked for contamination.

### Mice and *in vivo* metastatic melanoma studies

All animal studies were conducted following the guidelines and regulations of the Ethics Committee for Animal Experimentation of Federal University of São Paulo (UNIFESP). 6-8 Week-old healthy male C57BL/6 (Wild Type, WT) or NOD/Scid/IL-2rγnull (NSG) mice were obtained from CEDEME (Centro de Desenvolvimento de Modelos Experimentais para Medicina e Biologia - UNIFESP) and were housed in ventilated racks (ALESCO) in specific pathogen-free conditions (SPF). Mice (n=5, per group) were intravenously challenged with 5 × 10^5^ (for WT) or 5 × 10^4^ (for NSG) syngeneic B16F10 viable cells in 0.1 mL of RPMI medium without fetal bovine serum (FBS), and treated on the next day with intraperitoneal (i.p.) doses of 300 μg (10 mg/kg) of C36L1 peptide, for 5 consecutive days, or with control vehicle (PBS). After 14 days, mice were euthanized and lungs were harvested and assessed for metastatic colonization. The number of metastatic lesions was quantified using a stereo microscope (Magnification, ×4) (Nikon, Tokyo).

### Tissue paraffin immunofluorescence

Deparaffinization and antigen retrieval were performed at low pH and 96 °C cycles for 1h using a PT-link system (Dako). Tissues were washed 3 times in PBS and permeabilised in 0.1% triton-X100 for 2 minutes. Tissues were washed in PBS and blocked with 8% goat serum in PBS for 2 hours at RT in a humid chamber, then incubated with the following primary antibodies: anti-iNOS (Abcam), anti-Arg1 (Bioss) and anti F4/80 (Biolegend), anti-CD8 (Biolegend), anti-granzyme B (Abcam) overnight at 4 °C in a humid chamber. Next, tissues were washed 3x in PBS, and incubated with the secondary antibodies goat anti-Rat IgG Alexa Fluor 594 (Abcam), goat anti-rat IgG Alexa Fluor 488 (Abcam), donkey antirabbit IgG DyLight 488 (Biolegend) and 10 μg/mL Hoechst 33342 (Sigma-Aldrich) for 2 hours at RT in the dark. Tissues were washed in PBS and slides were mounted with mounting solution (Dako) and sealed with nail polish. Images were acquired using an Axio Observer Light Microscope with the Apotome.2 (Zeiss). Metastatic melanoma lesions were gated by generating a region of interest (ROI) and positive fluorescence was calculated among ROIs using the NIS-Elements Advanced Research 4.0 software (Nikon, Tokyo).

### Flow cytometry analysis of intratumoral dendritic cells and lymphocytes

C57BL/6 mice were previous challenged with metastatic B16F10 melanoma cells and treated with C36L1 peptide or control vehicle (PBS) as described above. For dendritic cells analysis, lungs were harvested and digested with collagenase A 1mg/mL for 1 hour at 37 °C, filtered in a 70 μm pore cell strainer and 5 lungs from each group were pooled together. Pan DCs were purified using the mouse Pan Dendritic Cell Isolation Kit according to manufacturer’s instructions (Miltenyi Biotec, Bergisch Gladbach, Germany). Both enriched CD11c^+^ and CD11c^-^ cell fractions were used for DCs and lymphocyte analysis, respectively. Cells were resuspended in PBS + 5% BSA and blocked for 20 minutes at 4 °C with TruStain anti-FcX (BD Pharmingen, Clone 2.4G2) and probed with the following conjugated antibodies for 1 h at RT: anti-CD11c (V450), anti-CD86 (PE-Cy7), anti-MHC-II (V500), anti-CD197 (PERCP-CY5.5), for DC analysis and anti-CD3 (PE) in combination with anti-CD4 (FITC), anti-CD8 (FITC) and anti-NK1.1 (FITC) for lymphocyte analysis. All antibodies were purchased from BD Pharmingen (Franklin Lakes, NJ, EUA). Cells were washed three times and immediately analysed by flow cytometry using the FACSCanto II (Becton Dickinson, San Jose, CA, USA). Acquired data was analysed using the FlowJo V10 software (TreeStar Inc., Ashland, OR, USA). Experiments were performed in triplicates.

### Flow cytometry analysis of splenic T regulatory cells and macrophages

Fresh spleens were obtained from C57BL/6 mice bearing melanoma lung metastasis and treated with C36L1 peptide or control vehicle (PBS). Splenocytes were isolated as described before, 5 spleens were pooled together from each group and probed with the following conjugated antibodies: anti-CD4 (FITC) and anti-Foxp3 (APC) for lymphocyte analysis, anti-F4/80 (FITC), anti-CD86 (PE-Cy7) and anti-CD40 (APC) for macrophage analysis. All antibodies were purchased from BD Pharmingen (Franklin Lakes, NJ, USA). Data from splenic Treg analysis represent three *in vivo* experiments. Data from macrophages represent flow cytometry absolute values of five pooled spleens per group from one *in vivo* experiment.

### TGF-β ELISA assay

CD11c^+^DCs (1x10^5^) were purified from lymphoid tissues of C36L1 treated mice and control vehicle (PBS) using the mouse Pan Dendritic Cell Isolation Kit according to manufacturer’s instructions (Miltenyi Biotec, Bergisch Gladbach, Germany). DCs were seeded in 12-well plates and cultured for 48h at 37°C. The supernatant was collected and TGF-β levels were measured using the Mouse-TGF-beta ELISA Set (BD, OptEIA ™) detection kit according to the manufacturer’s instruction. Data represent mean ± s.e.m of [TGF-] values from three biological replicates.

### Tumour conditioned medium preparation

1x10^6^ B16F10 melanoma cells were cultured in 175 cm^2^ culture flasks and in complete RPMI-1640. When cells reached 70% of confluence, the medium was harvested, centrifuged at 2000 rpm and filtered using a 0.22 μm pore filter, concentrated using StrataClean Resin (Agilent Technologies), and immunoblotted for MIF detection, or stored at -20 °C for further use in functional assays. Alternatively, to increase the concentration of tumour-secreted factors, B16F10 cells were sub-cultured in TCM and fresh media (v/v).

### Generation of bone marrow derived macrophages and myeloid dendritic cells

For generation of primary macrophages and myeloid DCs, bone marrow cells were isolated from the femurs of C57BL/6 mice in cold MAC buffer (Ca^2+^, Mg^2+^ free PBS + 2 mM EDTA + 0.5% BSA), centrifuged at 1200 rpm for 10 min, re-suspended in 5 mL RBC Lysis Buffer (1X, BD Pharm Lyse) and incubated for 5 min at RT. Reaction was terminated in PBS and cells were centrifuged at 1200 rpm for 10 min at RT. Cells were re-suspended in 5 mL of MAC buffer and carefully added in the top of 5 mL of Histopaque solution (Sigma-Aldrich) in 15 mL tubes and centrifuged at 1200 rpm, 25 min at 15°C without brake and 1 acceleration. The monocyte-enriched fraction was collected in a new tube and washed in PBS. Monocytes were further incubated with M-CSF-1 (10 ng/mL) in complete DMEM media (Thermo Fisher) to generate macrophages (13), or GM-CSF (50ng/mL) plus IL-4 (25ng/mL) in complete RPMI to generate myeloid DCs, mainly characterized by CD11b^high^CD11c^high^ (17, 33). In the 3^rd^ day of monocyte differentiation, medium was replaced with new cytokine enriched fresh medium and cultured for 3 more days. To generate macrophage conditioned media (MCM) for the experiment described in Figure 6, macrophages were treated with TCM, MIF (200ng/mL) or left untreated, in the presence or absence of C36L1 peptide (200 μM) for 72h, and further incubated in serum free medium for 48h. After 48h, the medium was harvested, centrifuged, filtered and stored at -20°C.

### CD8^+^ T cells isolation from naïve splenocytes

Fresh spleens were obtained from healthy C57BL/6 mice and placed into cell strainer immersed in MAC buffer on a Petri dish. Using the plunger end of the syringe, spleens were mashed through the cell strainer and splenocytes were resuspended in 15 mL of MAC buffer and centrifuged at 300 g for 5-10 minutes at room temperature. Supernatant was discarded and the pellet resuspended in 5 ml of red blood cell lysis buffer (Sigma Aldrich; Missouri, USA) for 5 minutes with occasional shaking. The reaction was terminated by diluting the Lysis Buffer with 10 mL of 1X PBS. Cells were washed in PBS and quantified using Trypan Blue. The negative CD8a^+^ T Cell Isolation Kit (Miltenyi Biotec, Bergisch Gladbach, Germany) was used to isolate CD8^+^ T lymphocytes from spleens of C57BL/6 mice following the manufacturer’s instructions.

### Flow cytometry analysis of primary macrophages and DCs

For flow cytometry analysis of primary macrophages and myeloid DCs, cells were detached using PBS/EDTA 10mM, scraped and collected in 1.5 mL Eppendorf tubes, centrifuged at 2000 rpm for 5 min, washed 1x in PBS and re-suspended in blocking buffer (PBS/BSA 1% plus TruStain fcX anti-mouse CD16/32 (Biolegend)) on ice for 30 min. Cells were washed 1x in PBS and incubated with the primary antibodies for 40 min on ice, following the manufacture’s instructions for each antibody. Cells were stained with the following antibodies: anti-F4/80 (FITC), anti-CD86 (PE.Cy7), CD40 (APC-A), anti-CD11c (APC), anti-CD11b (FITC), anti-MHC-II (Percp-Cy5.5), anti-CD80 (PE-Cy7), anti-CD86 (PE), all purchased from Biolegend, anti-CD197 and Pacific Blue from Becton Dickinson. After incubation with antibodies, cells were washed 3x in PBS, fixed in paraformaldehyde (PFA) 2% and acquired using the Attune Acoustic Focusing Cytometer (Applied Biosystems, Foster, CA, USA). Analysis was performed using FLowJo v10 software (Tree Star, Ashland, OR, USA). All antibody-panels were designed without channel colour overlay and antimouse/anti-rat compensation beads were used for control compensation procedures following the manufacturer’s instructions (BD, Franklin Lakes, NJ, USA).

### Immunofluorescence and confocal microscopy

Sterile glass coverslips were seeded at the bottom of 24 well plates and 1x10^5^ B16F10, primary macrophages and DCs were seeded in 500 μL of their respective culture media and cultured overnight. For detection of MIF in B16F10 cells, media was removed and cells were washed in PBS and fixed in 2% paraformaldehyde (PFA) for 15 min at RT, washed and permeabilised in 0.1% PBS-Triton X100 for 15 min at RT and blocked in blocking buffer) solution (PBS-BSA 2% plus 5% goat serum) for 30 min. Primary rabbit anti-MIF antibody (Abcam) was added for 1hour at 4 °C. Three consecutive washes in PBS were performed and slides were incubated with secondary antibody solution (anti-rabbit IgG Alexa Fluor 488 (Abcam) and 10 μg/mL of Hoechst 33342) was added for 1hour at RT and protected from light. Slides were mounted using DAKO mounting solution and sealed with nail polish. For assessment of CD74 expression and interaction with C36L1 in primary macrophages and myeloid DCs, cells were initially incubated with biotinylated C36L1 peptide for 30 min at 300 μM, further washed and blocked with blocking buffer for 15 min at 4 °C and fixed in PFA 2% for 15 min. Cells were incubated with primary mouse-anti-CD74 (Abcam) for 1h at 4 °C and further incubated with secondary solution: streptavidin-Alexa Fluor 594 (Red) (LifeTechnology) plus anti-mouse IgG Alexa Fluor 488 (Abcam) (Green) and 10 μg/mL of Hoechst 33342 (Sigma-Aldrich) for 1 hour at RT. Coverslips were washed and placed on the top of glass slides with DAKO mounting solution and sealed with nail polish. For immunofluorescence and confocal analysis, images were acquired using an Axio Observer Fluorescence Microscope with the Apotome.2 (Zeiss) or a confocal Zeiss Microscope, respectively. The merge region between red and green channels (shown in white in Figure 5) was quantified using the automated threshold analysis tool of the ImageJ analysis software. At least three different fields per image were quantified.

### Primary macrophages and myeloid dendritic cells culture assays

Primary macrophages and myeloid DCs were generated as described above. 5x10^5^ cells were seeded in 12-well plates in complete fresh media and 200 μM of C36L1 peptide was added to the cultures for at least 6 h prior to the addition of B16F10 TCM or 200 ng/mL of recombinant MIF (R&D System, Minneapolis, MN, USA). Cells were incubated at 37 °C for 72 hours and further used in FACs analysis for phenotyping or functional assays. All experiments were performed in triplicates.

### Dendritic cells stimulation for CD8+ T cell activation assays

Primary myeloid DCs were generated from C57BL/6 mice as described above, and 2.5x10^4^ cells were seeded in 48-well plates in quintuplicates and incubated for 24 hours at 37 °C. C36L1 (200 μM) peptide was added to the cultures 6 h prior to incubation with recombinant MIF at 200 ng/mL for 72 hours. After incubation, cultures were treated with 200 μM of the tyrosinase-related protein 1 (TYRP-1) peptide (NDPIFVLLH) as a MHC class I specific melanoma antigen. Previous naïve CD8+ T cells purified from C57BL/6 splenocytes, as described above, were pre-activated for 24 hours at 37 °C using 30U/mL of IL-2 and anti-CD3/CD28 dynabeads following the manufacturer’s instructions (Thermo Fisher). Next, primary myeloid DCs were washed and incubated with 2.5x10^5^ CD8^+^ T cells, in the presence of 30U/mL of IL-12 (PeproTech, London, UK). Co-culture was maintained for 5 days and primed CD8^+^ T cells were harvested and co-cultured with 2.5x10^3^ B16F10 melanoma cells (10:1) for 72 hours. After 72 hours, CD8+ T cells were harvested from the cultures and the remaining viable B16F10 cells were quantified with a Neubauer chamber using the Trypan Blue dead cells exclusion stain and using the MTT colorimetric based assay.

### B16F10 proliferation assay with macrophage conditioned media

To obtain different macrophage conditioned media, primary macrophages were cultured in the following conditions for 72 hours: (1) alone, (2) in the presence of tumour conditioned medium (TCM) or with recombinant MIF (200ng/mL) and (3) pre-incubated for 6 hours with C36L1 peptide (200 μM) followed by TCM or MIF (200ng/mL) incubation. Next, the medium was removed and macrophages were further cultured with serum free medium for 48 hours to produce macrophage conditioned media corresponding to the different conditions (MCM1, MCM2 and MCM3). MCM was harvested from the different macrophage culture conditions, filtered through 0.45 μM and added to 2x10 B16F10 melanoma cells plated in 96-well plates stained with CFSE (Thermo Fisher). B16F10 melanoma cells were cultured with the different MCMs for 72 hours. Next, B16F10 cells were harvested from wells, stained with propidium iodide (10 μg/mL) and the total number of viable (PI^-^) and proliferating cells (CFSE^-^) was quantified by flow cytometry acquiring fixed volumes of cell suspension using an Attune Flow Cytometer.

### Quantitative real-time PCR (qPCR) experiments

Total RNA from primary macrophages previously stimulated with C36L1 (200 μM) for 6 h and recombinant MIF (200 ng/mL) for 72 h was isolated using the RNeasy^®^ Mini Kit (Qiagen, Hilden, Germany). cDNA was prepared from 100ng RNA per sample, and qPCR was performed using gene-specific QuantiTect Assay primers (Qiagen) following the manufacturer’s instructions. qPCR reactions were performed using FIREPol^®^ EvaGreen^®^ qPCR Mix Plus ROX (Solis Biodyne, Tartu, Estonia) in a MaxQuant system. The following primers were used: TGF-P (Mm_Tgfb1_1_SG, Qiagen), IL-10 (Mm_ IL10_1_SG, Qiagen), PD-L1 (Mm_Pdcd1Ig1 _1_SG, Qiagen), Arginase-1 (Mm_ Arg1_1_SG, Qiagen), IL-6 (Mm_Il6_1_SG, Qiagen), GAPDH (Mm_Gapdh_3_SG, Qiagen). Relative expression levels were normalized to *Gapdh* expression according to the formula <2^(*C*_tgene of interest_ – *C*_tgapdh_)^ (13), and displayed as fold change units.

### Protein extraction and immunoblotting

Primary macrophages and myeloid DCs (5x10^5^ per well) were obtained as described above, serum starved for 24 hours, treated with C36L1 (200 μM) for 6 hours (or left untreated) and stimulated with recombinant MIF (200 ng/mL) for 5, 10 or 20 min to detect phosphorylation of AKT (10 and 5 min for primary macrophages and myeloid DCs, respectively) and ERK1/2 (20 and 5 min for primary macrophages and myeloid DCs, respectively). Cells were lysed in RIPA buffer (150 mmol/L NaCl, 10 mmol/L Tris-HCl pH 7.2, 0.1% SDS, 1% Triton X-100, 5 mmol/L EDTA) supplemented with a protease and phosphatase inhibitors solution cocktail (Invitrogen), 1 mmol/L phenyl methyl sulfonyl fluoride and 0.2 mmol/L Na_3_VO_4_, and protein concentration was quantified using BCA assay (Thermo Fisher). Protein lysates were separated by electrophoresis and immunoblotting analyses were performed for: total AKT (CST: 9272S), total p44/42 MAPK (ERK1/2) (CST: #4695), phospho-AKT (Ser473) (CST:4060S), phospho-ERK1/2 (Thr202/Tyr204) (CST: #4370). GAPDH (Sigma), was used as protein loading control. To verify the presence of MIF in the tumour conditioned medium (TCM), TCM was filtered with 0.45-μM filter and concentrated using StrataClean Resin (Agilent Technologies), and immunoblotted for MIF (Abcam). HRP-conjugated secondary antibodies (Cell Signaling Technology, Beverly, MA, USA) were used followed by incubation with the ECL substrate (Pierce). Images were acquired using a transillumminator Alliance 9.7 (Uvitec, Cambridge UK). Phosphorylation ratios were quantified using ImageJ gels’ algorithm, normalized to untreated control lanes.

### Peptide/Protein binding prediction

The computational modelling platform Pepsite 2.0 (Russel-Lab) (34) was used to predict the binding probability of peptides to mouse MIF (PDB: 1MFI, chain B) and mouse CD74 (PDB: 1IIE, chain B) proteins. Results are displayed as p values, where p ≤ 0.05 values are the statistically significant binding predictions. iCDR peptide was used as a negative peptide control. Binding probability was calculated using the interval 0.01 < p < 0.05, where p = 0.01 represents 100% of binding probability and p > 0.05 represents 0% of binding probability.

### C36L1 preparation and molecular dynamics

We obtained the 3D structure of C36L1 by performing de novo structure prediction in Pep-Fold3 web-server. In order to obtain a set of equilibrated conformations to perform molecular docking experiments, we carried out a molecular dynamics (MD) simulation on GROMACS 5.1 using CHARMM36 force field. We set up the simulation system on CHARMM-GUI web-server, with TIP3P water molecules and Cl^-^ counter ions to charge balancing. After 5000 steps of steepest descent energy minimization, a NVT equilibration dynamics of 25 ps and then a NPT production MD of 100 ns was carried out. We clustered the MD trajectory to obtain a diverse conformational population to perform molecular docking. All MD frames were fitted to the reference structure and clustered with GROMOS method by using GROMACS 5.1, with a backbone root-mean-squared deviation (RMSD) cutoff of 5.0 Å for C36L1 resulting in 8 different clusters. Thus, the centre structure of each cluster was then used in docking simulations.

### CD74 normal mode calculations and generation of low-energy conformations

The CD74 structure 1IIE (35) (residues from 118 to 176) was used to perform normal modes analysis (NMA) by using CHARMM c41b1, and CHARMM36 force filed using DIMB module. A distance dependent dielectric constant was employed to treat the electrostatic shielding from solvation. We computed the 5 lowest-frequency normal modes (generally the most collective ones) as directional constraint to generate low-energy conformers along the mode trajectory using the VMOD algorithm in CHARMM, as previously described (36, 37). The restraints were applied only on Cα atoms and the energy was computed for all atoms. The structures were displaced from –3.0 Å to +3.0 Å using steps of 0.1 Å, resulting in 61 intermediate energy relaxed structures along each mode.

### Molecular docking

Molecular docking simulations were performed with ATTRACT using iATTRACT algorithm. The general strategy is to depict both aspects that govern binding dynamics between two macromolecules at the same time: conformational selection and induced fit. In the conformational selection model, proteins and ligands are both described as an ensemble of conformations in equilibrium in solution being the active conformers more probable to interact. Induced fit, however, occurs after the initial interaction, when protein side chains undergo structural changes to better adapt to the ligand. To do this we provide several conformations of both receptor and ligand (ensemble docking) and the docking software combines simultaneous interface flexibility and rigid body optimizations during docking energy minimization. The best 50 solutions were written for each combination. We used BINANA 1.2 to investigate the specific molecular basis guiding the interaction between CD74 and C36L1.

### Chemiluminescent Dot blot binding assay

Peptide C36L1 binding to CD74 was determined by chemiluminescent (CL) dot-blotting carried out as previously described (24). Briefly, 25 nmoles of C36L1 and the irrelevant CDR peptide control (iCDR) and vehicle (0.025% DMSO in dH_2_O) were carefully dotted in 10 μL of dH_2_O on nitrocellulose membranes, and blocked with 1% TBS-Tween (TBS-T) in 5% BSA for 1h at 4 °C. Membranes were washed in TBS-T and incubated with 25 nM of recombinant CD74 (Abcam) in the presence of protease inhibitors in PBS/BSA overnight at 4 °C, using a shaker. After incubation, membranes were washed three times in TBS-T for 5 min, and incubated with primary mouse anti-CD74 (Abcam, Cambridge, UK). Membranes are washed and incubated with secondary anti-mouse IgG-HRP (CST) for 2 hours at 37 °C. Immunoreactivity was determined using the ECL Western Blotting Substrate (Pierce™) and signal was detected in a transilluminator Alliance 9.7 (Uvitec, Cambridge UK).

### Statistics

All statistic tests were performed using the GraphPad Prism 5.0 software (San Diego, CA). Statistical differences between experimental and control group were calculated using the Student’s *t*-test. All experiments were performed in triplicates. Significant differences are indicated by *p < 0.05, **p < 0.01 and ***p < 0.001.′

## RESULTS

### The Anti-metastatic effect of the C36L1 peptide requires the immune system

We have previously shown that intraperitoneal injections of the anti-tumour CDR peptide C36L1 significantly decrease pulmonary melanoma metastasis in a syngeneic model (24, 25). In addition, bone marrow derived myeloid pro-inflammatory dendritic cells (DCs) displayed equivalent anti-tumour effect when tumor antigen-primed DCs were pre-treated with C36L1 *ex vivo* and adoptively transferred to mice bearing lung melanoma metastasis (24). These findings suggest that the anti-tumour effects induced by C36L1 *in vivo* may result from the peptide ability to stimulate the host immune response. To further investigate the mechanism of action of C36L1, we treated immunocompetent C57BL/6 and immunodeficient NOD/Scid/IL-2rγnull mice bearing melanoma lung metastasis with C36L1 peptide or control vehicle (Figure 1A). We observed that C36L1 significantly decreased lung metastasis in immunocompetent mice but not in immunodeficient mice (Figure 1B). These findings confirm that C36L1 anti-tumour effect is driven by its ability to stimulate the immune response against metastatic melanoma.

**Figure 1.**
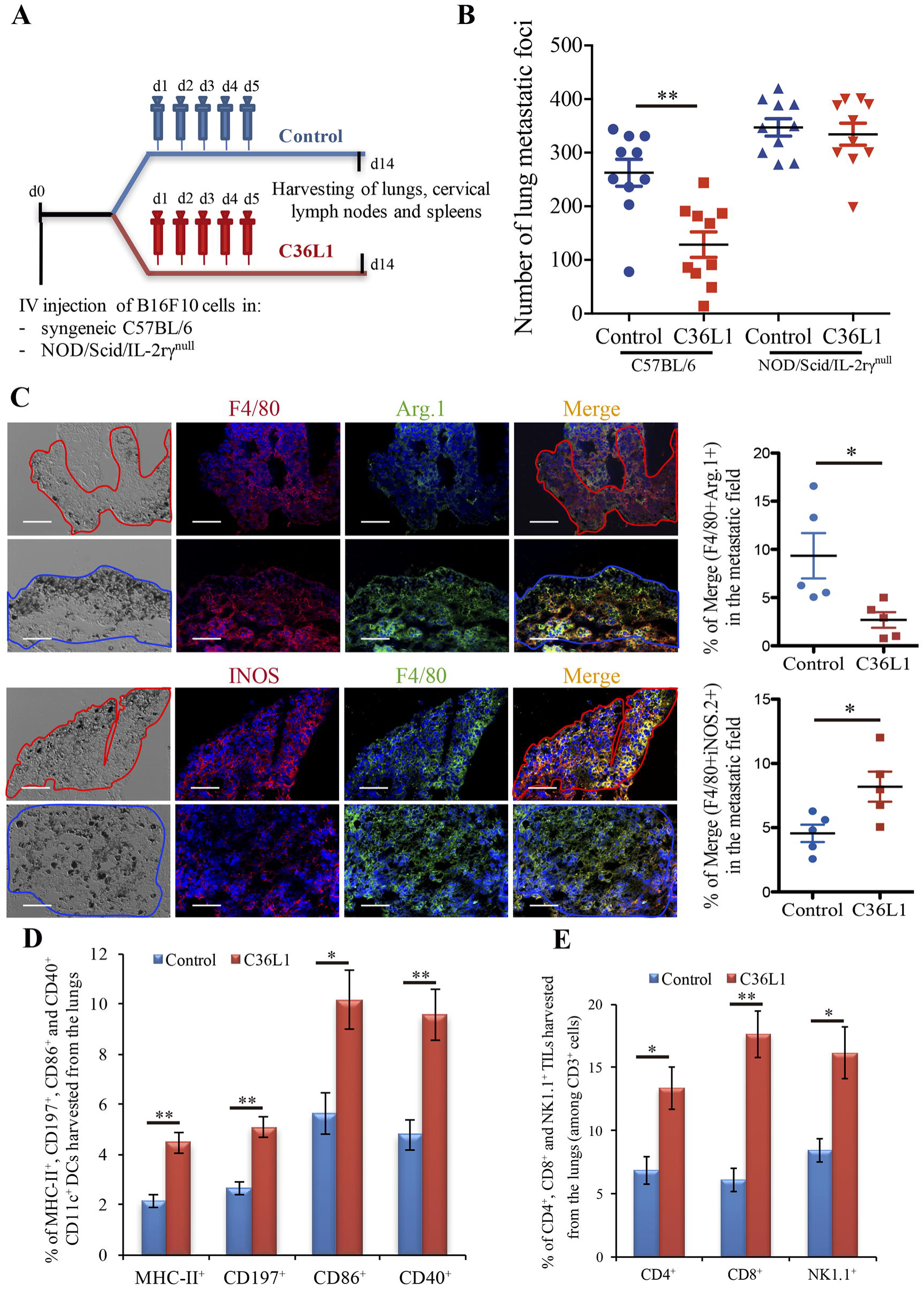
The anti-metastatic effect of the C36L1 peptide depends on the immune system. **(A)** Metastatic melanoma model and therapeutic strategy using C36L1 peptide and control vehicle (PBS). At end point, lungs, cervical lymph nodes and spleens are harvested. **(B)** Number of metastatic foci in immunocompetent (Wild Type, WT) and immunodeficient (NOD/Scid/IL-2rγnull, NSG) mice treated with control vehicle (PBS) or C36L1 peptide. *n* = 10 mice per group (two combined experiments). Values are expressed as means ± s.e.m., and were analysed using a two-tailed unpaired *t*-test. ** *p*=0.001. Graph combines two independent experiments. **(C)** Left, Immunofluorescent staining and quantification of F4/80^^+^^Arg.1^+^ M2-like and F4/80^+^iNOS^+^ M1-like macrophages in lung metastasis from C36L1 and control vehicle treated mice. Melanoma lung metastatic area appears in dark/brown colour in brightfield images. Right, Graphs show quantification of positive F4/80^+^Arg1^+^ (\**p*=0.028) and F4/80^+^iNOS^+^ (\**p*=0.02) stainings. Nuclei were counterstained with Hoechst 33342 (Blue). *N* = 5 mice per group; at least five fields assessed per sample. Values are expressed as means ± s.e.m., and were analysed using a two-tailed unpaired *t*-test. Blue and red lines indicate the tumour area in C36L1 and control vehicle treated mice, respectively. Scale bars: 50 μm. **(D)** Flow Cytometry quantification of activation markers MHC-II (\*\**p*=0.003), CD197 (\*\**p*=0.002), CD86 (\**p*=0.019), and CD40 (\*\**p*=0.007) expressed in CD11c^+^ DCs isolated from lungs of C36L1 and control vehicle treated mice. Data represent quantification of four independent experiments. Values are expressed as means ± s.e.m, and were analysed using a two-tailed, unpaired *t*-test. **(E)** Quantification of CD4^+^ (\**p*=0.03), CD8^+^ (\*\**p*=0.005) and NK1.1^+^ (\**p*=0.02) cells among CD3^+^ cells in lung metastatic lesions from C36L1 and control vehicle treated mice. Bar graphs combine three independent *in vivo* experiments with 5 pooled lungs per group for each experiment. Values represent means ± s.e.m., and were analysed using a two-tailed unpaired *t*-test.

### C36L1 restores macrophages and DCs immunogenic functions in metastatic melanoma

Macrophages and DCs are vital for activating effector T cells and shaping the immune response against cancer (7). In solid tumours, including melanomas, macrophages and DCs are suppressed by the tumour and lose their ability to activate and support the immune response against cancer (12, 17). Tumour associated macrophages (TAMs) often acquire an M2-like phenotype that hampers the anti-tumour immune response and supports tumour growth, metastasis and resistance to therapies (12-14, 38). Similarly, intratumoral DCs often acquire a tolerogenic phenotype and lose their ability to activate effector T cells (17, 39, 40). Thus, effective anti-cancer immunotherapies must reverse the tumour immunosuppressive environment and restore the immunogenic functions of macrophages and DCs. In this respect, we found that C36L1 is able to re-polarise M2-like (F4/80^+^ Arg1^+^) tumour associated macrophages into M1-like (F4/80^+^ iNOS^+^) pro-inflammatory and anti-tumorigenic macrophages (Figure 1C). In addition, increased levels of M1-like macrophages were also observed in the spleens of C36L1 treated mice, compared to control treated mice (Supplementary Figure 1A). The number of activated intratumoral DCs (CD11c^+^, MHC-II^+^, CD197^+^, CD40^+^ and CD86^+^) in metastatic lungs from C36L1 treated mice was significantly increased compared to control treated mice (Figure 1D). C36L1 treatment decreased the secretion of the immunosuppressive cytokine TGF-β by CD11c^+^ DCs from lymphoid organs (spleens and cervical lymph nodes) (Supplementary Figure 1B). These findings suggest that C36L1 re-polarises and re-activates macrophages and DCs’ immunogenic and anti-tumorigenic functions in metastatic melanoma.

### C36L1 increases the level of effector T cells in the TME

Tumour specific antigen presentation by DCs and macrophages to effector T cells is a crucial step for the generation of an effective immune response against cancer, and increased infiltration of effector T cells in tumours is a good prognostic marker (4, 5). Since treatment with C36L1 decreases melanoma pulmonary metastasis and increases the numbers of pro-inflammatory macrophages and DCs, we asked whether C36L1 increases effector T cell infiltration in metastatic tumours. We found that, indeed, C36L1 significantly increased the levels of CD4^+^ T cells from 6.86% to 13.35%, CD8^+^ cytotoxic T cells from 6.11% to 17.6%, and NK1.1^+^ natural killer cells from 8.44% to 16.13%, in lung metastatic melanoma (Figure 1E and Supplementary Figure 2A). In lymphoid organs, tolerogenic DCs are responsible for inducing Foxp3^+^ Tregs differentiation by secreting TGF-β. Since C36L1 treatment decreases TGF-β production by DCs (Supplementary Figure 1C), we evaluated whether CD4+Foxp3+ Tregs were also reduced in lymphoid organs upon C36L1 treatment. Flow cytometry analysis of mice splenocytes revealed a highly significant decrease in the percentage of CD4^+^Foxp3^+^ Tregs from 59.6% to 1.39% following C36L1 treatment (Supplementary Figure 2B). Together, these findings indicate that C36L1 restores DCs and macrophages immunogenic functions, increases effector T cell infiltration in metastatic tumours and inhibits immunosuppressive regulatory T cells.

### C36L1 inhibits the suppressive effects of tumour-secreted factors in macrophages

Tumour educated macrophages exhibit an M2-like phenotype and support cancer progression in several ways, including the direct support of cancer cell proliferation (17). To further understand how C36L1 affects macrophage function, we cultured metastatic B16F10 melanoma cells with conditioned media from tumour educated macrophages (macrophages previously exposed to tumour conditioned media) in the presence or absence of C36L1. As expected, melanoma cells exposed to tumour educated macrophages showed a significant increase in proliferation. Addition of C36L1 abrogated this macrophage - driven tumour cell proliferation (Figure 2 and Supplementary Figure 2C). These results show that macrophages exposed to tumour conditioned media acquire pro-tumorigenic functions and this can be inhibited by C36L1 peptide. These findings suggest that C36L1 must interfere with a tumour secreted factor (or its receptor) that regulates macrophage function.

**Figure 2:**
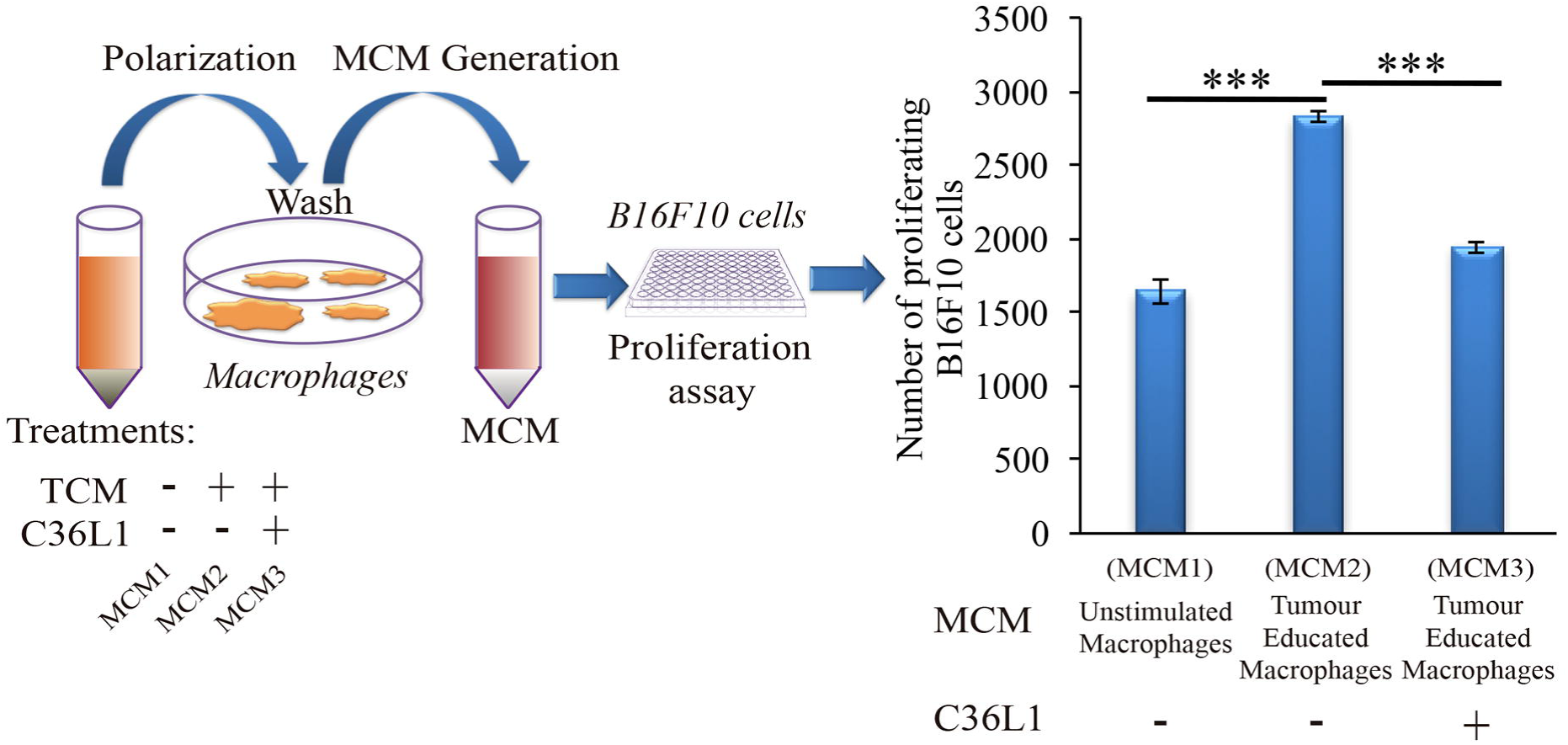
C36L1 counteracts the pro-tumorigenic activity of macrophages induced by melanoma derived factors. **Left:** Schematics describing the workflow of the tumour cell proliferation assay. Tumour cells are exposed to either conditioned media from: untreated macrophages (MCM1), macrophages exposed to tumour conditioned media (TCM) from metastatic melanoma B16F10 cells (MCM2), or macrophages exposed to C36L1 peptide + TCM from B16F10 cells (MCM3). Next, MCM generated from these three conditions were added into B16F10 melanoma cells and the number of live proliferating cells was quantified by flow cytometry after 72h. **Right:** Bar graph represents average of three independent experiments. Values represent means ± s.e.m. and data were analysed using a two-tailed unpaired *t*-test. *** p<0.001.

### C36L1 binds to MIF receptor, CD74

C36L1 is a linear and flexible CDR-based peptide. Linear peptides are likely to adopt a few stable conformations and interactive possibilities to different relevant targets (41). Previous studies have shown that stromal and melanoma cells express high levels of MIF, supporting melanoma growth and modulating immune cells in late-stage melanoma (29, 30, 42-46). Dendritic cells and macrophages both express MIF’s main receptor, CD74 (47). Thus, we hypothesize that C36L1 could interfere with MIF signalling on macrophages and dendritic cells. In agreement with previous studies, we observed that B16F10 metastatic melanoma cells express and secrete high levels of MIF *in vitro,* (Figure 3A, B), and that MIF is highly expressed in small and large lung metastatic melanoma lesions (Figure 3C).

**Figure 3:**
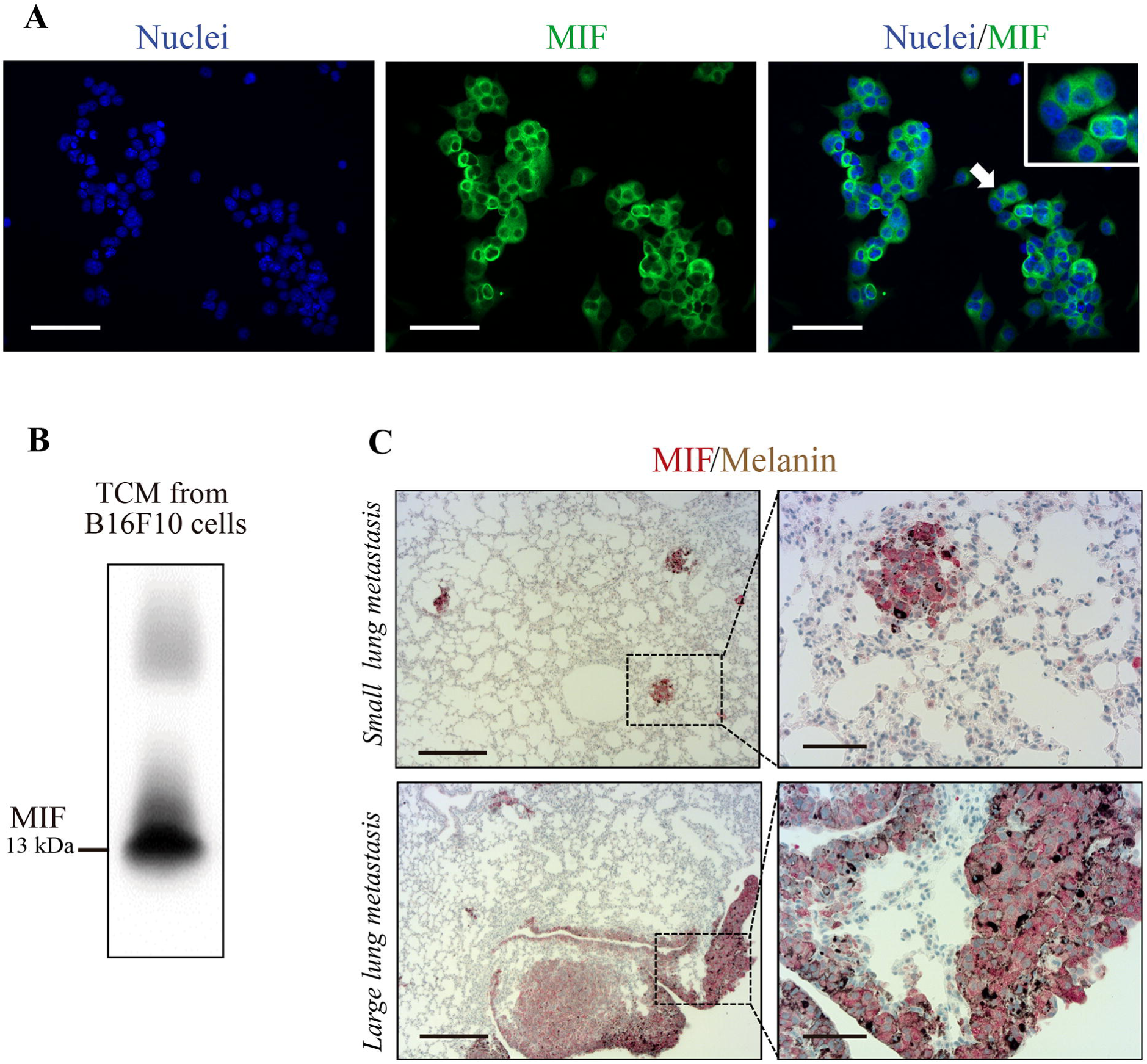
MIF is secreted by B16F10 metastatic melanoma cells and is highly expressed in lung metastatic lesions. **(A)** Immunofluorescent staining of B16F10 cells stained for MIF (green) and nuclei (blue). Scale bars: 50 μm. **(B)** Immunoblotting analysis of B16F10 tumour conditioned media (TCM) detecting secreted MIF. **(C)** Immunohistochemical staining of lung melanoma metastasis showing MIF (in red) in small and large lesions. Dark brown areas are metastatic foci of melanoma cells. Scale bars: 200 μm (Left) and 50 μm (Right).

A pilot study addressing the binding probability of C36L1 to MIF and its receptor CD74 was carried out using the computational modelling prediction of peptide-binding sites to protein surfaces and the Pepsite 2.0 algorithm (34). This *in silico* approach predicted a statistically significant binding of C36L1 to mouse CD74 B chain (PDB: 1IIE) protein (p < 0.001), and a potential binding to mouse MIF B chain (PDB: 1MFI) protein (p = 0.04) (Figure 4A). No interaction with either CD74 or MIF was predicted for an irrelevant control CDR peptide (iCDR - CE48-H2), which was previously observed to have no effect on metastatic melanoma proliferation *in vitro* and progression *in vivo* (25) (Figure 4A and Supplementary Figure 3A). We used the Pepsite 2.0 algorithm to identify the amino acid residues involved in the interaction of C36L1 to CD74, and found that the peptide is predicted to interact with Tyr (118), Arg (179) and His (180) residues from the B chain of the murine/human CD74 protein, highlighted in red (Supplementary Figure 3B). Interestingly, Mesa-Romero et al., have recently described that some of these residues (highlighted in green) are also critical for the interaction of MIF with the CD74 antagonist (RTL-1000) (48). The *in silico* predicted interaction of C36L1 with CD74 was further confirmed in a dot-blot binding assay using both immobilized C36L1 and iCDR peptides against recombinant murine CD74 protein (Figure 4B). These results suggest that C36L1 could act as an antagonist of MIF, since its interaction occurs on critical binding sites used by MIF to interact with CD74.

**Figure 4:**
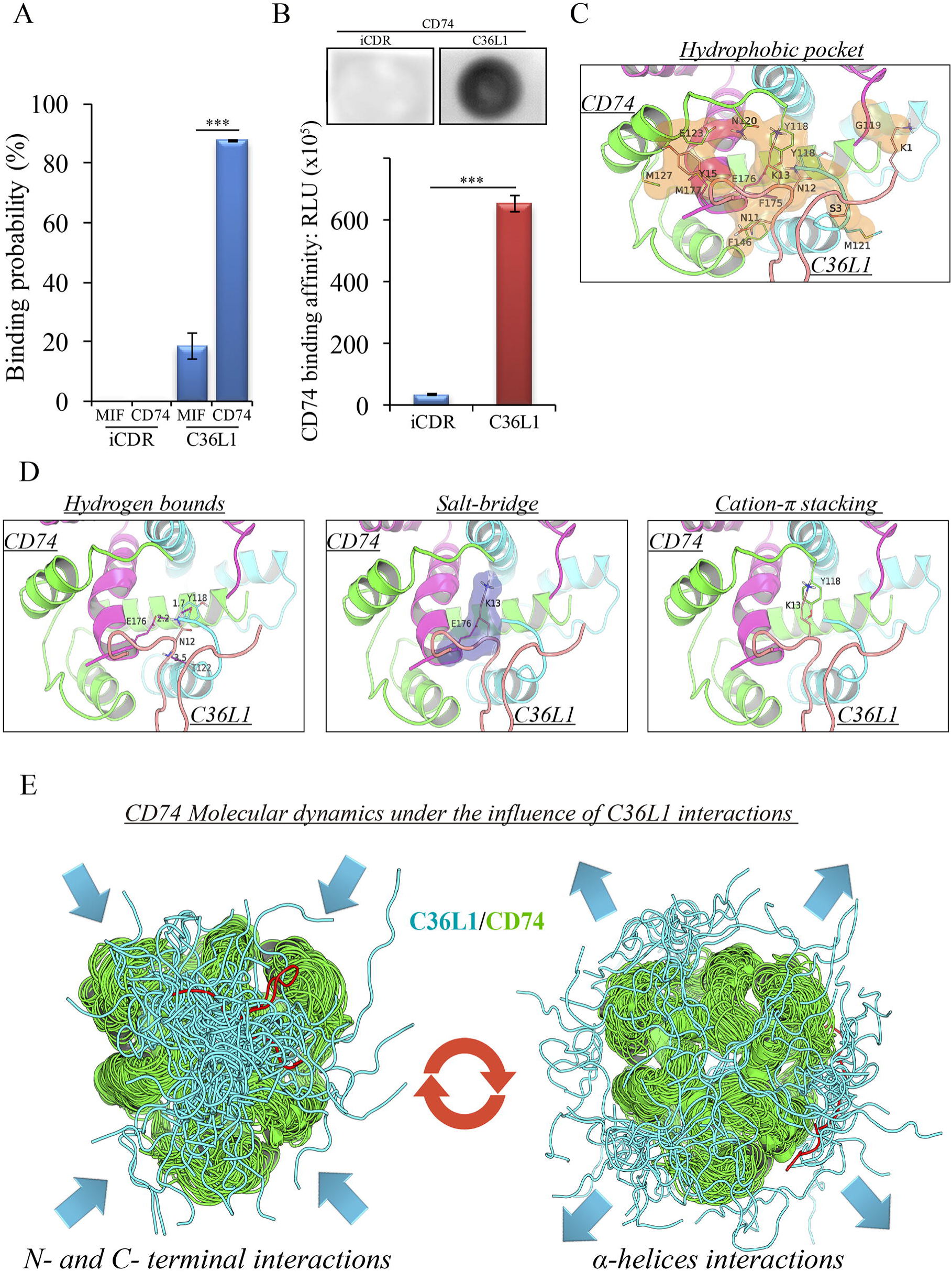
Binding prediction and molecular docking of C36L1 dynamic interactions to MIF and its receptor CD74. **(A)** Binding probability of C36L1 peptide and irrelevant peptide (iCDR) to MIF and its receptor CD74 calculated using Pepsite algorithm (*** p<0.001). **(B)** Dot-blot binding assay for C36L1 and iCDR peptides to mouse recombinant CD74. Bar graph represents mean of RLU in dot area quantified using ImageJ software (***p<0.001). **(C)** Hydrophobic pocket (orange) formed by CD74 and C36L1 partners characterized by carbon-carbon interactions above a 4Å distance cut-off. **(D)** Electrostatic interactions between CD74 and C36L1 peptide: hydrogen bonds formed between partners. Donor-acceptor distances are described; salt bridge formed involving K13; cation-n stacking between tyrosine residues of chain A of CD74 and C36L1. CD74 chains A, B and C are coloured in green, cyan and magenta, respectively. C36L1 is coloured in yellow. **(E)** Overlap of highest and lowest free energy results for C36L1 (cyan) in complex with CD74 (green). **Left**: Overlap of the lowest free energy 50 poses showing major concentration of C36L1 peptide at the CD74 N- and C-terminal interface. Lowest peptide free energy pose highlighted in red. **Right:** Overlap of the lowest free energy 50 poses, where C36L1 visits other regions of CD74, including the external region of the α-helices. Lowest peptide free energy highlighted in red.

To further investigate this, we performed a molecular docking study between C36L1 and CD74 protein. Docking calculations resulted in 122,000 different poses of which the worst 1% were discarded for presenting outliers’ energy values. The average energy of remaining structures was 60.8 kcal/mol and more than 95% of them presented thermodynamically favourable binding energies (Supplementary Figure 3C). The best solution occurred between C36L1 cluster 5 centroid and a CD74 structure with large opening (2.7 Å from reference) along normal mode 10, which shows an open-close motion. This pose presented -192.6 kcal/mol as free energy of binding, and is depicted in Supplementary Figure 3D. The key interaction elements observed in this complex were analysed using BINANA algorithm. Hydrophobic contacts forming an extended pocket along the interface of all CD74 subunits were observed (Figure 4C). Stronger interactions were also observed: three critical hydrogen bonds, one salt-bridge and one cation-π stacking interaction between CD74 and C36L1 peptide (Table 1 and Figure 4D). Interestingly, C36L1 cluster 5 centroid appears in 30 of top 50 best poses suggesting that this peptide conformation is likely to be privileged to bind CD74. Moreover, structures with large displacements along mode 10 of CD74 are more frequent; the worst ranked structures were less displaced. C36L1 interacts better with CD74 as it moves according to normal mode 10, whereas once CD74 returns to the relaxed conformation, the peptide binding affinity decreases and the complex dissociates. Furthernore, the overlap of 50 best solutions showed a putative preferred binding region of C36L1 to the interface formed between N- and C-terminal portions of CD74 monomers. This binding site is corroborated by the observation of C36L1 main binding to CD74’ α-helices, only in the worst solutions. In Figure 4E, blue arrows indicate spatial distribution of C36L1 (blue) over CD74 altered structures (green) and the best and worst poses of C36L1 are shown in red. A video representing the consequences of this dynamic interaction between C36L1 and CD74 tertiary structure is shown in Supplementary Video 1.

### C36L1 binds to CD74 on macrophages and DCs and disrupts downstream signalling

CD74 is a transmembrane protein mainly expressed in APCs and associated with the MHC II intracellular trafficking. CD74 is the main receptor for MIF in macrophages and DCs, and MIF binding to CD74 leads to immunosuppression of macrophages, activation of myeloid derived suppressor cells (MDSCs), suppression of natural killer (NK) cells and inhibition of T cell activation (29, 43, 47, 49-51). Thus, we evaluated whether C36L1 peptide (as predicted in the *in silico* approach) physiologically binds to CD74 receptor on macrophages and DCs.

To address these interactions, primary bone marrow derived macrophages and DCs were incubated with biotinylated C36L1 probed with streptavidin-PE (Red), and stained for CD74 (green). We observed that C36L1 binds to CD74 in both macrophages and DCs (Figure 5A). CD74 can be expressed intracellularly and at the plasma membrane. Using confocal microscopy, we observed that C36L1 co-localizes with CD74 both intracellularly and at the cell membrane (Figure 5B). MIF interaction with CD74 receptor activates different cell signalling pathways, including the PI3K/AKT and the MAPK signalling pathways (47, 49, 52). In agreement with this, we observed that recombinant MIF induces the phosphorylation of AKT (S473) and ERK (Thr202/Tyr204) in both primary macrophages and DCs (Figure 4C). However, pre-incubation of macrophages and DCs with C36L1 inhibited MIF induced AKT and ERK downstream signalling on macrophages and DCs. These findings show that C36L1 binds to CD74 on macrophages and DCs and disrupts MIF-CD74 signalling on these cells.

**Figure 5:**
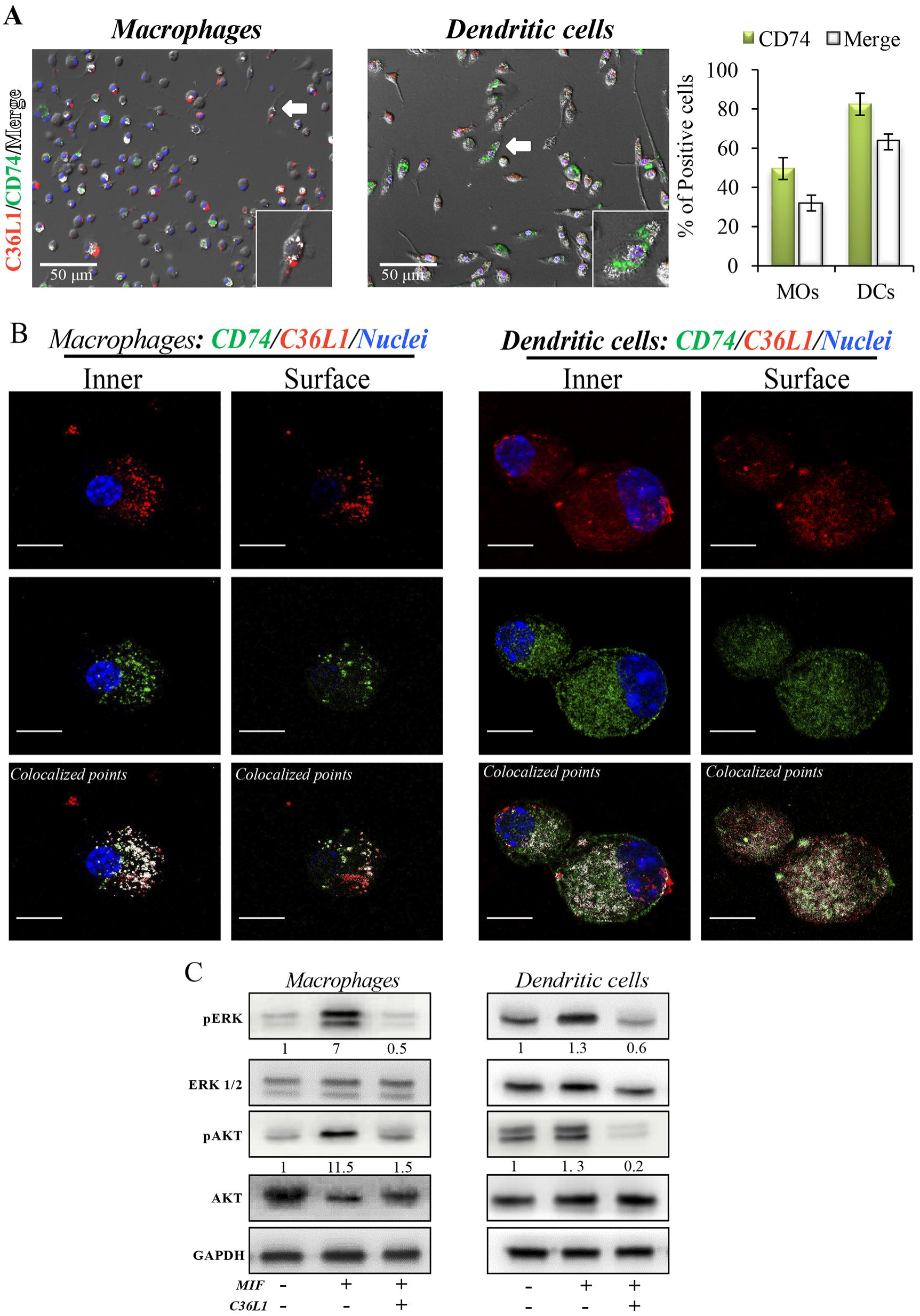
C36L1 interacts with CD74 in both macrophages and dendritic cells, and inhibits MIF/CD74 signalling. **(A)** Immunofluorescent staining of C36L1 (red), CD74 (green), nuclei (blue) in primary macrophages (MOs) and dendritic cells (DCs). CD74 interactions with C36L1 were quantified using automated analysis in ImageJ. Arrows indicate merged channels depicted in white. Four fields per slide were quantified. Scale bars: 50 μm. **(B)** Representative fluorescent confocal microscopy images showing colocalisation of C36L1 peptide (Red) and CD74 (green) in the intracellular and surface focal plane of both primary macrophages (left) and DCs (right). Co-localized points were detected using ImageJ colocalization algorithm, depicted in white. Scale bars: 10 μ,m. **(C)** Immunoblotting analysis of phosphorylated AKT and ERK1/2 on primary macrophages and DCs previously treated with C36L1 (200 μg/mL) or left untreated, and further treated with recombinant MIF (200 ng/mL).

### C36L1 inhibits MIF induced suppression of macrophages and DCs and restores their immunogenic and anti-tumorigenic functions

To further understand the mechanism of action of C36L1 on macrophages, we evaluated the immunosuppressive and tumour supporting functions of macrophages exposed to MIF in the presence or absence of C36L1. Macrophages exposed to MIF supported the proliferation of melanoma cells (similar to what we observed when we exposed macrophages to tumour conditioned media (TCM) in Figure 2). C36L1 treatment abolished this MIF-induced pro-tumorigenic function of macrophages (Figure 6A). C36L1 also significantly decreased the expression of the immunosuppressive factors TGF-β, IL-10, Arginase-1, PD-L1 and IL-6 by macrophages exposed to MIF (Figure 6B).

**Figure 6:**
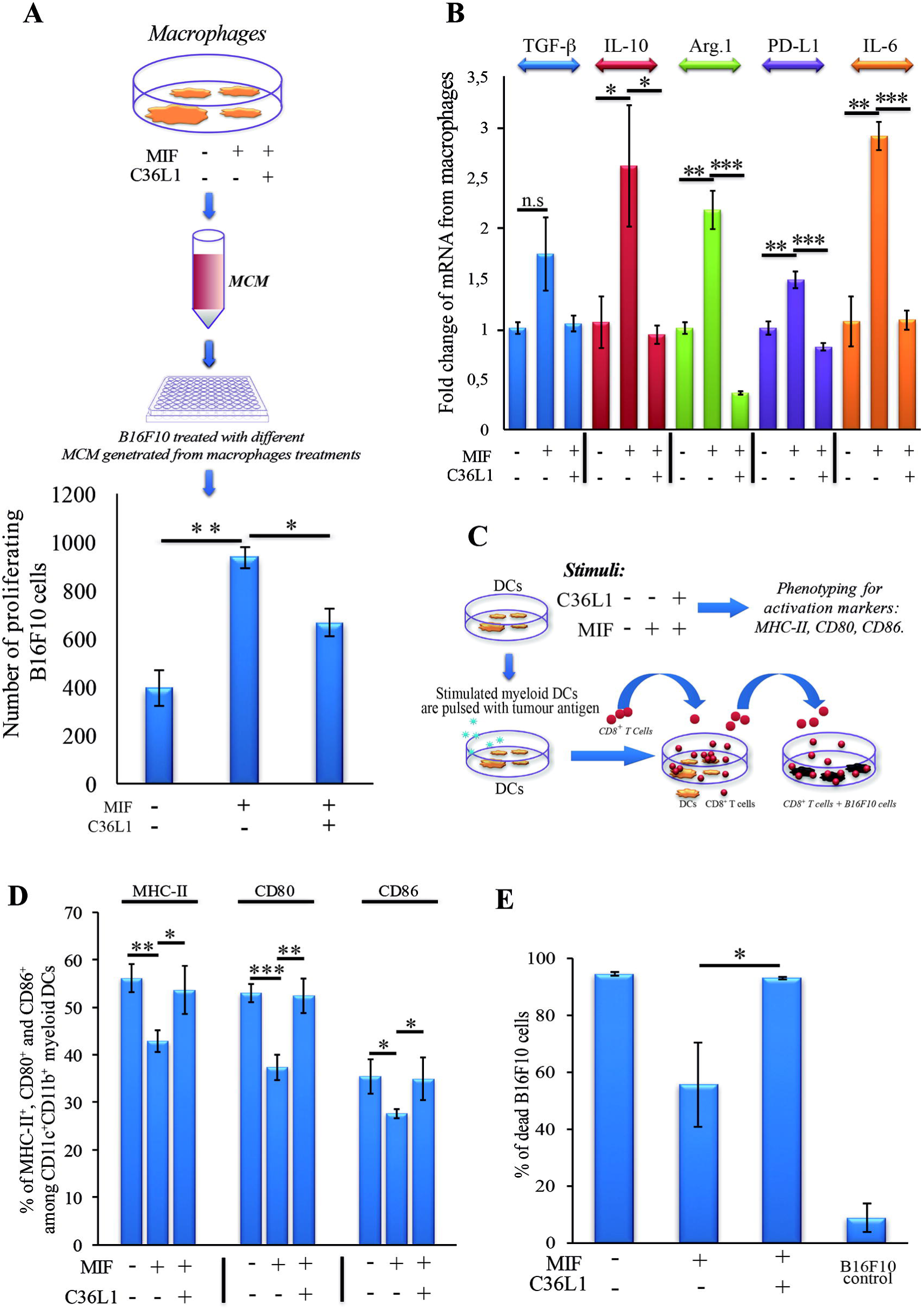
C36L1 blocks MIF induced immunosuppressive effect on macrophages and dendritic cells. **(A) Top:** Schematics describing the workflow of the tumour cell proliferation assay. B16F10 metastatic melanoma cells are exposed to conditioned media from: untreated macrophages, macrophages exposed to MIF (200 ng/mL) or macrophages exposed to C36L1 (200 μg/mL) + MIF (200ng/mL). The number of live proliferating B16F10 cells was quantified by flow cytometry after 72h. **Bottom:** Bar graph represents average of three independent experiments, mean ± s.e.m. Data were analysed using a twotailed unpaired *t*-test. *** p<0.001. **(B)** C36L1 blocks MIF induced immunosuppressive effect on primary macrophages. mRNA levels of *TGF-P* (n.s = 0.058), IL-10 (\**p*=0.049, *p*=0.042), Arg.1 (\*\**p*=0.002, ***p<0.001), PD-L1 (\*\**p*=0.0049, ***p<0.001), and IL-6 (**0.0015, ***p<0.001) from macrophages exposed to recombinant MIF in the presence or absence of C36L1 peptide. Experiments were performed in triplicates. Values represent mean ± s.e.m and were analysed using a two-tailed unpaired *t*-test. **(C)** Schematics describing the different conditions in which DCs were cultured and then used to activate T cells. Primary DCs were incubated with MIF (200 ng/mL) in the presence or absence of C36L1 peptide and activation markers were quantified by flow cytometry. These DCs were further pulsed with a melanoma antigen peptide and incubated with syngeneic purified CD8+ T cells. Next, T cells were harvested and incubated with melanoma B16F10 cells at a ratio of 10/1 CD8^+^ T cells/ B16F10 tumour cell. **(D):** Quantification of MHC-II (*p*=0.01), CD80 (*p*<0.001) and CD86 (*p*=0.02) activation markers in DCs performed by flow cytometry. Bar graph represents mean ± s.e.m, from three independent experiments. Data were analysed using a two-tailed unpaired *t*-test. **(E)** Bar graph showing the quantification of dead B16F10 cells after incubation with CD8^+^ T cells. Best of three independent experiments is shown, mean ± s.e.m from biological triplicates, one-tailed unpaired *t*-test (\**p*=0.032).

To understand the mechanism of action of C36L1 on DCs, we evaluated the expression levels of DC activation markers as well as DCs ability to activate cytotoxic T cells in the presence or absence of MIF and C36L1 (Figure 6C). Treatment of primary myeloid DCs with MIF significantly decreased the levels of the maturation and co-stimulatory markers CD86, CD80 and MHC-II. Treatment with C36L1 peptide counteracted the immunosuppressive effect of MIF on DCs (Figure 6D). DCs ability to activate cytotoxic T cell killing function was also significantly impaired by MIF but rescued by C36L1 treatment (Figure 6C, E, Supplementary Figure 4). All together, these results provide functional evidence that C36L1 restores DCs and macrophages immunogenic and anti-tumorigenic functions by interfering with the MIF/CD74 immunosuppressive signalling axis.

## DISCUSSION

Cutaneous melanomas are common in the Western hemisphere causing the majority (75%) of deaths related to skin cancer (53). The incidence rate of melanoma increases faster than for any other cancer (52). At very-early stages, melanomas can be surgically removed and the 5-year survival rate of melanoma is 98%. However, melanoma can metastasize to distant organs including lungs, liver, bones and brain, and the 5-year survival rate of patients with metastatic melanoma drastically decreases to 15-20% (1, 2).Treatment with immune checkpoint inhibitors has significantly increased the 5-year survival rate of melanoma patients (1, 55), but the number of non-responders is still high, with the lack of response being currently intensively investigated. Mutations of gene families of cytokines, chemokine levels, mesenchymal transition, E-cadherin and other proteins expressed in tumours are being studied (56). Understanding and targeting the immunosuppressive tumour microenvironment to restore an anti-tumour immune response is an area of great interest (7, 29, 57, 58). Therefore, understanding the mechanisms by which metastatic melanoma suppresses antitumour immunity could further contribute to the development of new combinatorial agents that restore the immune response against metastatic melanoma.

Synthetic peptides based on Immunoglobulin-CDR sequences have shown promising antitumour properties, and some of these peptides display immune stimulatory functions (22, 24-26). The C36 V_L_ CDR1 peptide (C36L1) was initially identified as an anti-melanoma agent (25) that inhibits melanoma cells survival and proliferation (24).

In this study, we uncover the mechanism by which C36L1 restores an effective immune response against metastatic melanoma. Specifically, we found that C36L1 is able to repolarise M2-like immunosuppressive tumour associated macrophages into immunogenic and anti-tumorigenic M1-like macrophages. C36L1 also promotes the activation and immunogenicity of DCs. C36L1 driven activation of the innate immune system leads to the inhibition of immunosuppressive Tregs, the activation of effector T cells and subsequently to the killing of metastatic melanoma cells. Mechanistically, we found that C36L1 binds to the MIF receptor CD74 on macrophages and DCs, thereby inhibiting MIF immunosuppressive effect on these innate immune cells, and shifting the balance from an immunosuppressive tumour microenvironment into a pro-inflammatory immunogenic environment in which the anti-tumour immune response is reinvigorated.

Tumours, including melanomas secrete factors that inhibit the immune system. Among these factors, MIF has been recently shown to have immunosuppressive activities, in many cancers, including glioblastoma, breast, pancreatic cancer and melanoma (29, 30, 49, 59-61). Thus, MIF is an emerging attractive target for immunotherapy. In pancreatic cancer, MIF is an important downstream regulator of fibrosis that culminates in the recruitment of TAMs favouring metastasis (21). In a similar way, metastatic uveal melanoma cells secrete MIF to recreate the eye immune-privileged environment and to inhibit the immune response in the liver, favouring liver metastasis (42, 61). In cutaneous melanoma, MIF is produced by melanoma cells to support growth and induce immunosuppression (29, 51). However, the role of MIF in metastatic melanoma remains unclear. In glioblastoma, MIF can also induce pro-inflammatory functions, including M1-like macrophage polarization (59,63). Bevacizumab, a monoclonal antibody that targets VEGF may also interact and neutralize MIF in glioblastomas, inducing the polarisation of macrophages into the M2-like phenotype that contributes to therapy resistance (59). This dual and opposite effect of MIF on the immune response depends on the cytokine milieu in the tumour microenvironment and on the levels of MIF. In fact, very low or high concentrations of MIF are thought to suppress the immune response, while intermediate doses rather promote pro-inflammatory and antitumour effects (59).

Different drugs targeting MIF and its main receptor CD74 are in clinical development in many diseases, including cancer (31, 32, 48, 64-67). The MIF inhibitor 4-iPP is so far the only immunomodulatory agent described to be effective in melanoma, and has shown promising results in subcutaneous melanoma, associated to an increase in monocyte pro-inflammatory functions (30). The effect of blocking MIF-CD74 signalling in metastatic melanoma has not yet been investigated. Targeting CD74 seems to be a promising anti-cancer therapeutic strategy to disrupt MIF induced suppressive signalling effect on monocytes (31,49,67). The most well-characterized CD74 inhibitor is Milatuzumab, a monoclonal antibody approved for the treatment of chronic lymphocytic leukaemia with acceptable side effects in humans (67). However, in the field of drug discovery, peptide based approaches emerge with intrinsic advantages, compared to antibodies including their small size, lack of immunogenicity, high affinity, specificity to different targets, low toxicity, good tissue penetration and biocompatibility (22, 25, 26). Peptides can exert immunomodulatory functions and have been shown to neutralize immune checkpoint receptors in cancer (68-70). Indeed, linear peptides such as CDR peptides are flexible and likely to bind to different biologically relevant targets (41).

In this study, we found that C36L1 interaction with CD74 is sufficient to disturb MIF’s induced immunosuppression of macrophages and DCs (Figure 7). Our *in silico* studies show that the flexibility of this linear peptide allows its transient interaction with the CD74 receptor, disturbing its molecular dynamics in the cell membrane. C36L1-CD74 interaction seems to be crucial to disrupt CD74 interaction with MIF in both macrophages and DCs. The cell internalisation of CD74 conjugates is a well-known pharmacological characteristic of CD74 (50, 67), which has been recently explored as a drug-carrier strategy for the treatment of lymphomas and B cell malignancies (67). CD74 internalisation independent of MIF binding could impair the activation of downstream signalling (31, 71). In this regard, we found that C36L1 binds to CD74 at the cell membrane as well as in the intra-cellular space of macrophages and DCs. This suggests that C36L1 binding to CD74 may promote its cytosolic internalization making it unavailable for binding to MIF. MIF binding to CD74 activates the PI3K/AKT and MAPK signalling pathways, and both these pathways have been related to monocyte immunosuppression, and macrophage M2-like polarization (45, 47, 49, 52). In agreement with these studies, we found that C36L1 inhibits MIF induced AKT and ERK1/2 phosphorylation in both primary macrophages and DCs and restores their anti-tumorigenic and immunogenic functions (Figure 7).

**Figure 7:**
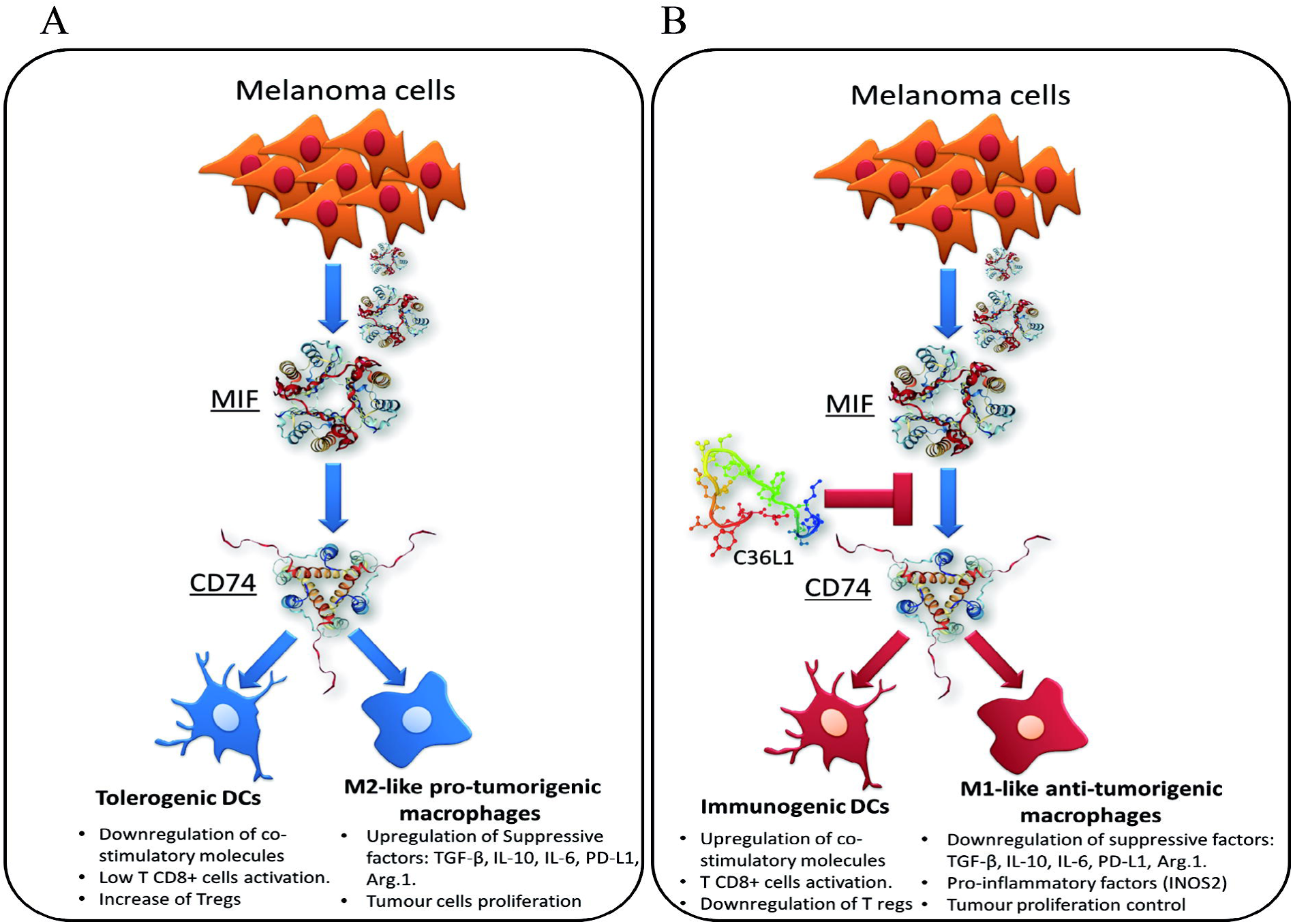
Scheme of the mechanism of action of the C36L1 peptide in macrophages and dendritic cells. C36L1 binds to MIF’s receptor CD74, thereby blocking its immunosuppressive effect on macrophages and DCs, restoring their anti-tumorigenic functions and their capacity to activate and support an effective immune response against metastatic melanoma.

In conclusion, our findings suggest that MIF is highly secreted in metastatic melanoma and is an important immunosuppressor of macrophages and DCs. Blocking MIF signalling through CD74 using the C36L1 Ig-CDR-based peptide restores the pro-inflammatory functions of macrophages and DCs thereby harnessing the immune response against metastatic melanoma. This study provides a rationale for further evaluation of CDR-based peptides as therapeutic agents to restore the ability of the innate immune system to start and shape an effective anticancer immune response.

## ETHICS STATEMENT

Animal experiments were carried out in accordance with the recommendations of the National Council for the Control of Animal Experimentation (CONCEA, Brazil), and performed according to the Ethics Committee of Federal University of São Paulo (CEUA N° 7588260915). Weight loss, lethargy and weakness that could result in inability to feed and drink, as well as infection with systemic signs of illness were considered as standard clinical symptoms that indicate deteriorating health conditions requiring euthanasia before the end of the experiment.

## ACKNOWLEDGMENTS

The authors acknowledge Ms Jennifer Adcott and Dr. Marco Marcello, from the Liverpool Centre for Cell Imaging (CCI) for providing additional support with the confocal microscopy studies.

## AUTHORS CONTRIBUTION STATEMENT

**CF** performed most of the experiments. **RA** performed *in vivo* experiments. **SM** assisted with flow cytometry experiments. **PR** performed the molecular docking and dynamics studies. **LI** helped with isolation of primary cells and with methodology development. **AS** assisted with tissue processing and IHC experiments. **NG** assisted with *in vivo* and flow cytometry procedures**. RC** assisted with molecular docking and dynamic analysis of C36L1/CD74 interaction model. **MS** assisted with methodology development and provided conceptual advice. **LP** assisted with the initial conception of Ig-CDR peptide biological functions. **LT** generated the peptide, assisted in its functional characterization and supervised the *in vivo* experiments. **AM** and **CF** designed experiments and wrote the manuscript. **AM** supervised the project. All authors helped with analysis and interpretation of results and approved the manuscript.

